# Silencing of an efflux pump coding gene decreases the efflux rate of pyrazinoic acid in *Mycobacterium smegmatis*

**DOI:** 10.1101/2021.10.29.466536

**Authors:** Stefany Quiñones-Garcia, Robert H. Gilman, Patricia Sheen, Mirko Zimic

## Abstract

**Background:** Tuberculosis (TB) is an infectious disease caused by *Mycobacterium tuberculosis* (MTB). The recommended treatment for TB is based on the use of first-line drugs, including pyrazinamide (PZA). PZA is also a drug used in the treatment of multidrug-resistant TB (MDR-TB) because of its main effect against the latent stage. The main cause of resistance to PZA is mutations in the *pnc*A gene, which compromise the activity of the encoded enzyme pyrazinamidase (PZAse), which hydrolyzes PZA into POA, the active antituberculosis molecule. The mechanism of action of PZA requires that POA is expelled from the bacterium by an efflux mechanism. After that, if the extracellular medium is sufficiently acidic, POA is protonated and returns to the cytosol, releasing the proton and repeating the cycle, resulting lethal to the bacteria. The efflux pump responsible for extruding the POA to the extracellular environment is not yet known. *Mycobacterium smegmatis* is naturally resistant to PZA and has a 900-fold faster POA efflux rate than MTB, and has the advantage to be a faster growing mycobacterium.

**Methods:** In the present study we have silenced the transcription of several genes encoding efflux pumps in *M. smegmatis* by CRISPRi (CRISPR interference). These genes (*MSMEG_0250, MSMEG_3815, MSMEG_0241, MSMEG_5046* and *MSMEG_0410*) were homologous to efflux pump genes in MTB. POA efflux rate was measured, and a quantitative Wayne’s test was performed after silencing each gene.

**Results:** Silencing of *MSMEG_0250*, resulted in approximately 5-fold decrease in the POA efflux rate in *M. smegmatis* (P<0.0001). None of the other silenced genes showed a notable decrease in POA efflux rate.

## INTRODUCTION

Tuberculosis (TB) is an infectious disease caused by *Mycobacterium tuberculosis* (MTB) and is one of the 10 leading causes of death worldwide^1,2^. In 2019, 10 000 000 cases of people were reported to have developed the disease with an estimated 1 200 000 deaths^1^.

Pulmonary TB infection is transmitted when a patient with active TB coughs or sneezes, expelling small particles in the form of aerosols that contain MTB and easily penetrate the respiratory tract^3,4^. The treatment recommended by the World Health Organization (WHO) to combat TB in adults is the use of first-line drugs (rifampicin, isoniazid, pyrazinamide, and ethambutol). ^5^. However, one of the main problems is multidrug-resistant TB (MDR-TB).^1^.

Pyrazinamide (PZA) is an important first-line drug used for the treatment of TB and the eradication of latent strains^6,7^. PZA enters MTB by passive diffusion, is hydrolyzed to its active form by the enzyme pyrazinamidase (PZAse), encoded by the pncA gene, to pyrazinoic acid (POA). By means of an efflux mechanism, POA is released into the extracellular environment, where it is protonated (HPOA), and then re-enters the bacterium assisted by a potential gradient, where it releases a proton (H^+^) acidifying the cytoplasm and repeating the cycle^8–10^.

Despite this understanding, the mechanism of action of PZA in MTB is not entirely clear. Neither the POA binding targets nor the efflux pumps responsible for transporting POA to the extracellular environment are known with certainty^6,11^. During the last decades, it has been identified that one of the main causes of resistance to PZA is mutations in the *pnc*A gene, which encodes PZAse, causing functional alterations^12–14^. Not only mutations in pncA, but also mutations in the promoter, which cause under-expression of PZAse, are associated with resistance to PZA^15^.

PZA-resistant strains that do not lose full PZAsa activity, suggest the existence of other complementary resistance mechanisms, such as possibly mutations in POA/PZA-binding target proteins or mutations in the efflux pumps that expel POA ^13,16–19^.

The mycobacterial cell wall, composed of mycolic acids, is a very efficient but permeable barrier that prevents the passage of some drugs into the intracellular environment of mycobacteria. However, the permeability characteristics do not explain the resistance to antituberculosis drugs^18,20,21^. Efflux pumps contribute to the expulsion of drugs to the extracellular environment, which decreases the concentration of drugs in the intracellular medium. Mutations in the efflux pump genes may limit the elimination of drugs from the intracellular medium and have been associated with bacterial resistance^18,22,23^. In the case of PZA, it has been observed that resistant strains have a low POA efflux rate, suggesting that POA efflux velocity could be a mechanism of study to identify efflux pumps related to the mechanisms of action and resistance to PZA^24,25^.

Recent studies have reported efflux pump genes that may be related to drug resistance, such as Rv1634 and *Rv1250*, which have been shown to decrease susceptibility to antibiotics when overexpressed in MTB^26,27^. Likewise, clinical strains of MTB that do not present mutations in PZAsa, but in the *Rv1183* and *Rv0202c* genes, which encode the transmembrane transporter proteins MmpL10 and Mmpl11 respectively, show resistance to PZA, while the *Rv0206c* gene, which encodes the MmpL3 protein, has been reported as a target for binding to antituberculosis drugs^28,29^.

*Mycobacterium smegmatis* has similar characteristics to *M. tuberculosis*, although it is not pathogenic, is faster growing and is naturally resistant to PZA. Therefore, *M. smegmatis* constitutes an appropriate model to study anti-tuberculosis drugs^30,31^.

In the present work, we silenced homologous MTB genes in *M. smegmatis, MSMEG_0250, MSMEG_3815, MSMEG_0241, MSMEG 0410*, and *MSMEG_5046* was performed by CRISPRi. The silenced strains were evaluated to measure the change in POA efflux velocity in *M. smegmatis*.

## METHODOLOGY

### Bacterial strain

All *M. smegmatis* strains generated in this study are derived from reference strain mc^2^155 and *Escherichia coli* from Nova Blue.

### Culture media

*E. coli* strains were grown on Luria-Bertani (LB) medium at 37°C. *M. smegmatis* strains were grown in 7H9 broth, and 7H10 media on plates supplemented with 0.2% de glycerol, 0.05% Tween 80, and 10% Oleic Albumin Dextrose Catalase (OADC). The antibiotics kanamycin (20 µg/µl) and carbenicillin (50 µg/µl) was used for *M. smegmatis* and *E. coli* cultures recombinant with pLJR962 plasmid.

### Growth of mycobacteria

*M. smegmatis* clones with and without the pLJR962 plasmids with the cloned gRNAs were cultured in 10 ml of 7H9 broth with kanamycin (20µg/µl) and carbenicillin (50µg/ml) at 37°C at 300 rpm for 3 - 5 days. From this culture was inoculated in a new 7H9 broth with kanamycin (20 µg/µl) and carbenicillin (50 µg/ml) at 37°C at 300 rpm for 18 hours, until reaching its logarithmic phase, OD_600nm_: 0.5 - 0.8. It was then diluted with a 7H9 medium until its OD_600nm_ was 0.1 - 0.2 to induce the CRISPRi system.

### Homologous genes of MTB efflux pumps in M. smegmatis

MTB genes reported to encode efflux pumps (*Rv1634, Rv0202c, Rv1183, Rv1250*, and *Rv0206c*), were used to identify the homologous genes in *M. smegmatis*, using the Basic Local Alignment Search Tool - NCBI (BLAST) ^32^, with an identity filter greater than 50%.

### Design and cloning of guide RNAs (RNAg)

Based on the homologous genes in *M. smegmatis* (*MSMEG_0250, MSMEG_3815*, and *MSMEG_0241*), an RNAg was designed for each according to the protocol described by Rock ^33,34^. Specifically, two RNAg were designed for each of the *MSMEG_5046* (gRNA1 and gRNA2) and *MSMEG_0410* (gRNA1 and gRNA2) genes.

The gRNAs were cloned into the plasmid pLJR962 having the dCas9_spy_ system, with the restriction enzyme BsmBI (New England) and into *E. coli* Nova blue. The plasmids were then extracted, and cloning was confirmed by sanger sequencing at MACROGEN (USA) and transformed by electroporation into *M. smegmatis*.

### Transcriptional Repression of Genes by CRISPRi

Co-expression of dCas9spy was induced with the gRNAs of each gene in *M. smegmatis* strains with 100 ng/mL ATc, which were incubated for 14 - 24 hours at 37°C, 300 rpm until reaching an OD_600nm_ greater than 1. In addition, as a control for each gene, we evaluated the recombinant bacteria with the CRISPRi system with no ATc added.

### mRNA quantification

Mycobacterium was induced and not induced with ATc were centrifugate mRNA quantification in a 15 mL RNase and DNase free tube at 12500 rpm for 20 min at 4°C, then the pellet was washed with TE buffer pH8 at 12500 rpm for 20 min at 4°C and resuspended in 1 mL of TRI Reagent (Zymo Research) in a screw-cap tube with 0.1 mm zirconia beads. Cells were lysed at 6.5m/s for 30 seconds, twice, with an interval of three minutes for each cycle in the Fast Prep equipment (MP Biomedicals) and RNA was extracted with Directzol RNA kit columns (Zymo Research). The extracted RNA was treated with Ambion DNaseI (Thermo Fisher Scientific), and a DNA strand was formed with the SuperScript IV retro transcriptase kit (Thermo Fisher Scientific). Quantification of the mRNA concentration of the gene of interest with respect to the constitutively expressed gene MSMEG_2758, which encodes sigma factor, was performed by qPCR-RT following the published ΔΔCt protocol^35^, with SYBR Green (Takara) and with specific primers for each of them^33^. Primers were designed in NCBI primer-blast software, placed after the gRNAs of each sequence.

### Measurement of POA efflux rate

POA efflux velocity of mycobacteria with the silenced genes was measured following the previously standardized protocol and POA quantification was normalized to Bradford (POA mg/protein) ^24,25^. Briefly, mycobacterial growth and transcriptional repression of genes were performed in 17 mL of 7H9 medium. These were centrifuged at 12500 rpm for 20 min at 4°C, the pellet was resuspended with 10mM citrate buffer pH7 until their OD_600nm_ of ATc-treated and untreated mycobacteria were adjusted equally. From the pellet suspension, 900 µL aliquots were made and tempered at 37°C for 20 min, then 100 µL of 10 mM PZA (Sigma Aldrich) was added and the mixture was incubated at 37°C at different time intervals 5, 10, 15, 15, 20, 20, 25 and 30 min. The reaction was stopped by centrifugation at 14800 rpm for 2 min, of which 500 µL of the supernatant, extracellular fraction, was saved for the Wayne reaction at 450nm. The control, time 0, was 450 µL of supernatant without incubation with 50 µL of 10mM PZA. Pellets from the reactions were resuspended with 500 µL of 10 mM citrate buffer at pH 7, cells were lysed at 100 °C for 20 min, and centrifuged at 10 000 rpm for 10 min. The supernatants were used to quantify lysate proteins by Bradford (Bio Rad).

### Data analysis

POA efflux rate was estimated by linear regression of POA concentration at times 0, 5, 10, 15, 15, 20, 25, and 30 min. To compare efflux velocities between CRISPRi-silenced and CRISPRi-non-silenced bacteria, multiple linear regression was used including an interaction term between time and an indicator variable (0/1) that accounts for whether CRISPRi was or was not induced. Analyses were performed at 5% significance using Stata statistical software version 14 (Stata Corp., College Station, TX).

## RESULTS

### MTB genes in *M. smegmatis*

Genes *Rv1634, Rv0202c, Rv1183, Rv1250*, and *Rv0206c*, homologous MTB genes in *M. smegmatis* with greater than 50% identity were identified, *MSMEG_3815, MSMEG_0241, MSMEG_0410, MSMEG_5046*, and *MSMEG_0250* respectively.

### Quantification of the level of transcriptional repression of CRISPRi-silenced genes

Three qPCR repeats of the strains ATc-treated and untreated CRISPRi strains, showed notable variations of the transcription level. 100-1000 folds changes were observed (Figure 1).

**Figure 1:**
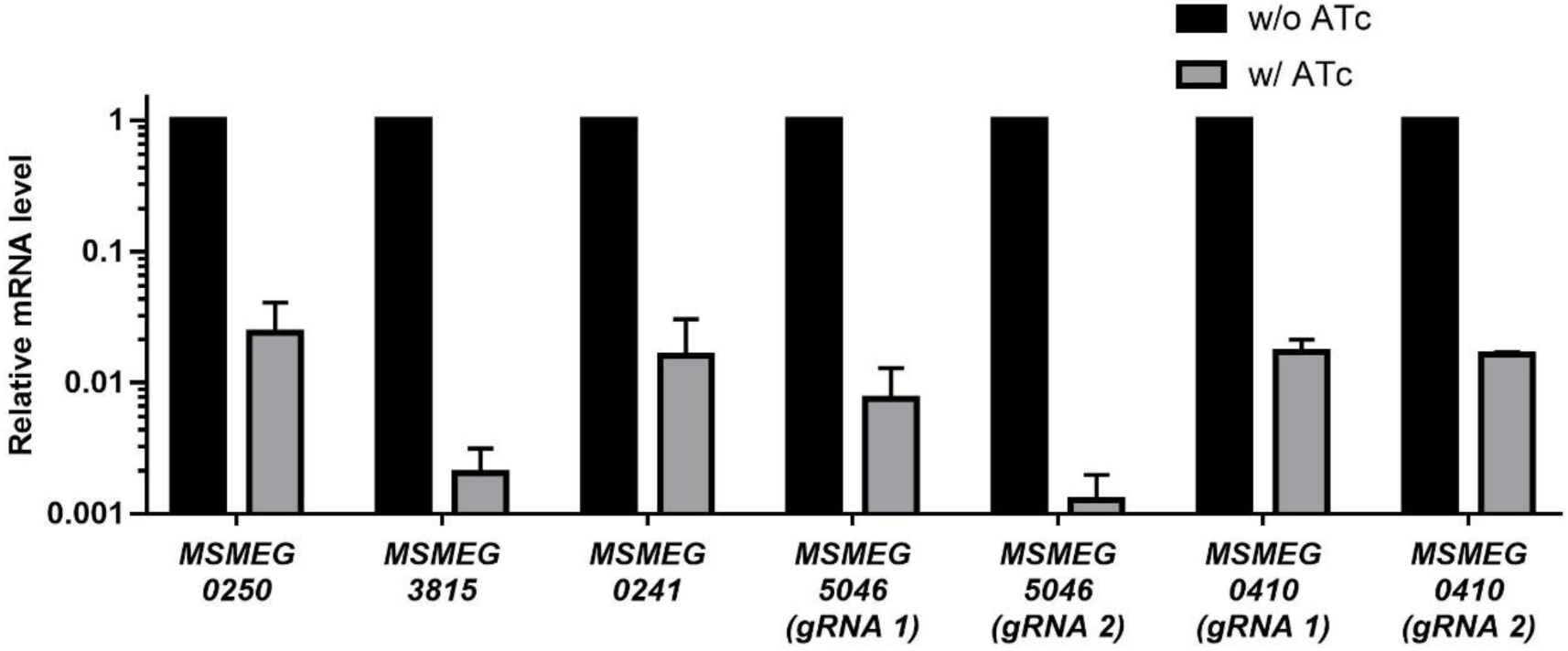
Plot of the difference of dCas9_sth1_ co-expression with MSMEG_0250, MSMEG_3815, MSMEG_0241, MSMEG_5046 (gRNA 1 and gRNA2) and MSMEG_0410 (gRNA 1 and gRNA2) genes by relative mRNA quantification, showing difference between bacteria with silenced, ATc treated, and unsilenced genes, without ATc. 95% confidence intervals are shown.

### Measurement of POA efflux rate in *M. smegmatis* strains with CRISPRi-silenced genes

The only gene that showed a noticeable and significant change in POA efflux velocity when CRISPRi was activated was the *MSMEG_0250* gene (p=0.000). The POA efflux rate without silencing (797 mM/s) was reduced 5-fold (162.8 mM/s). Although not significant, in *MSMEG_0241* and *MSMEG_5046* (gRNA 1) genes, a trend of a decrease in efflux velocity was observed in 1.4 and 1.6 folds respectively (Figure 2).

**Figure 2:**
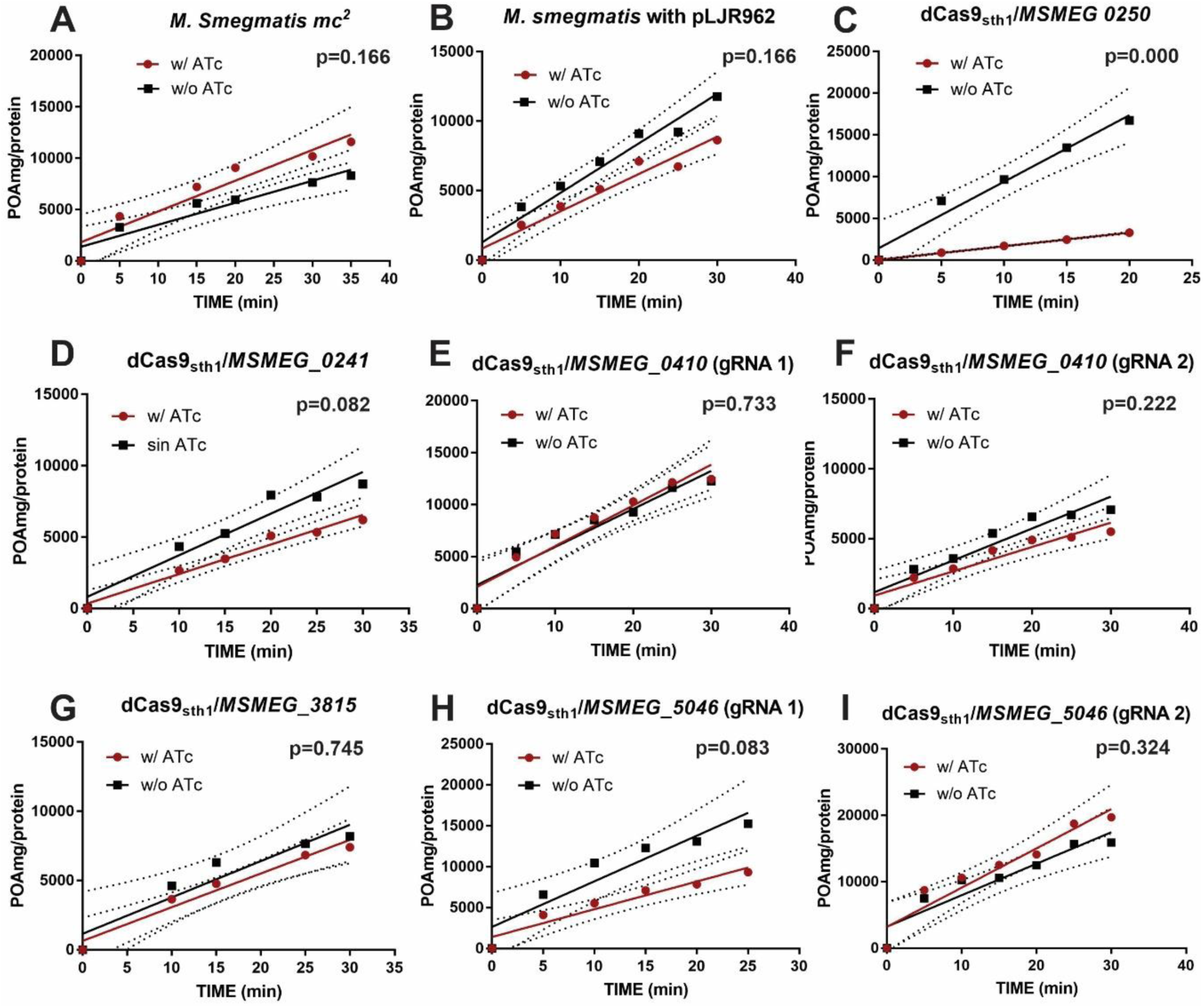
Measurement of POA efflux velocity in *M. smegmatis* when a gene is silenced with the CRISPRi system, induced with ATc and unsilenced, without ATc. (A) Control 1, *M. smegmatis*, no significant change in efflux velocity (P=0.166), (B) Control 2, *M. smegmatis* with plasmid pLJR962, no significant change in efflux velocity (P=0. 166), (C) Gene *MSMEG_0250*, shows significant difference in efflux velocity, when gene is silenced and unsilenced (P=0.000), (D) Gene *MSMEG_3815*, shows no significant difference in efflux velocity (P=0. 745), (E) Gene *MSMEG_5046* (gRNA 1), shows no significant difference in efflux velocity (P=0.083), (F) Gene *MSMEG_5046* (gRNA 2), shows no significant difference in efflux velocity (P=0.324), (G) Gene *MSMEG_0241*, shows no significant difference in efflux velocity (P=0. 082), (H) Gene *MSMEG_0410* (gRNA 1), shows no significant difference in efflux velocity (P=0.733) and (I) Gene *MSMEG_0410* (gRNA 2), shows no significant difference in efflux velocity (P=0.222). 95% confidence intervals are shown.

## DISCUSION

POA efflux velocity in *M. smegmatis* is markedly and significantly reduced when the *MSMEG_0250* gene is silenced by CRISPRi (p<0.001). Although not significantly, other genes also silenced are found to be associated with a trend of decreased POA efflux velocity.

Efflux pumps are responsible for generating the expulsion of drugs into the extracellular medium of the bacteria^6,9,10,24,25^. In previous works, we have observed that the rate of POA efflux into the extracellular medium has a wide range of variation, and thus possibly occurs through an active transport mechanism mediated by some efflux pumps (unpublished data). Typically, PZA-resistant strains tend to have a very low POA efflux rate; however, it has also been reported that PZA-resistant strains can have very high POA efflux rates. It appears that PZA-susceptible strains are associated with an intermediate range of POA efflux velocities^24^.

The *MSMEG_0250* gene, which encodes the MmpL3 protein, is an essential gene under in vitro conditions in mycobacteria and transporter of mycolic acid in the form of trehalose monomycolates (TMM) from the cytoplasm to the cell wall ^28,36,37^. Additionally, it is a complex bound to the TMM transport factor A (TtfA) protein, encoded by gene *0736* and to the *MSMEG_5308* gene. The first two complexes are exclusively responsible for TMM transport, while *MSMEG_5308*, upon Mmpl3 depletion, accumulates and stabilizes the Mmpl3/TtfA complex, preserving TMM transport and cell wall biosynthesis^38^.

The phenotypic evaluation shows that bacterial growth of *M. smegmatis* with the *MSMEG_0250* gene silenced by CRISPRi decreases on a 7H10 agar medium. This is not reproduced in 7H9 broth, where *M. smegmatis* grows in both silencing and non-silencing conditions. It is important to note that the liquid 7H9 medium allows mycobacteria to grow faster compared to the solid 7H10 medium^39^. This could explain the difference in growth in both media. In the relative quantification of transcripts (mRNA) of the *MSMEG_0250* gene, the decrease in expression under silencing conditions is partial with a Cq no greater than 30. It is possible that repression of the MSMEG_0250 gene is not sufficient to limit bacterial growth even though it is a highly vulnerable gene^33,40–42^. It is also possible that the partial loss of its function could be being compensated by other efflux pumps.

The vulnerability level of a gene is a measure of how essential it is. A highly vulnerable gene negatively affects the fitness of the bacterium and its growth^42^. The *MSMEG_0250* gene, in addition to having been reported as an essential gene, has recently been shown that both *MSMEG_0250* and its homolog *RV0206c* in MTB have a high vulnerability, and its silencing reduces bacterial growth ^42,43^. For this reason, we believe that the observed decrease in POA efflux velocity following *MSMEG_0250* gene silencing could potentially be due to loss of cell fitness. Further studies are needed to clarify this possibility.

Previous studies have shown that mutations in the *MSMEG_0250* gene decrease the hydrophobicity of the bacterial cell wall^28,44^. It is possible that a partial silencing of this gene could contribute to the change in the efflux velocity of POA, POA is a hydrophilic molecule^45^.

Also, recent studies based on the use of CRISPRi dCas9_sth1_ for the silencing of the gene homologous to *MSMEG_0250* in MTB, *Rv0206c* which encodes an efflux pump, have confirmed that this is a binding target for anti-TB drugs and can be used as an anti-TB drug discovery system^46,47^.

Inhibition of *MSMEG_0250* expression by CRISPRi could potentially affect the expression of more genes if they are associated with an operon^48^, especially if they are in the 5’ region of the gene of interest^33,49,50^. However, recent studies have shown that the *MSMEG_0250* gene does not share an operon with other genes, suggesting that the effect of silencing on POA efflux velocity would be a direct effect of the *MSMEG_0250* gene^42^.

During the evaluation of bacterial growth, no variation of bacterial growth was observed between mycobacteria silenced and non-silenced in the 7H9 culture medium. This confirms previous findings that ATc is not toxic to mycobacteria and would not be influencing the variation of POA efflux velocity ^25,51^.

ATc is a tetracycline derivative, an efficient inducer of the TetR repression promoter, which does not affect bacterial growth in *Mycobacterium*^33,51,52^. The optimal concentrations of ATc as an inducer for TetR in mycobacteria are from 50 to 100 ng/mL. At concentrations higher than 250 or 500 ng/mL, the inducer reduces its activity on the TetR promoter, decreasing the fold repression of silenced genes and increasing bacterial growth in essential genes_33,51,52_.

Although silencing of the *MSMEG_0410* and *MSMEG_5046* genes was performed with two different gRNA and different PAM sequences, which generated different repression folds, no significant variation of POA efflux velocity was detected (p>0.001). This strongly suggests that none of these efflux pumps would be involved in POA transport.

Although there are other systems for silencing mycobacterial genes and evaluating their effect on the rate of POA efflux, such as homologous recombination *knock-out*, it is important to recognize the advantage of using CRISPRi, since it is possible to control the level of repression of gene expression and thus avoid reaching extreme levels of total absence of expression^43,46,50,53^. Additionally, the CRISPRi system standardized by Rock *et al*. 2017, is easy and fast to manipulate. Thanks to the knowledge of PAM sequences and the different repression folds, it is simple to design a gRNA for a sequence of interest^33^.

Possible effects of *MSMEG_0250* gene silencing could be studied at the transcriptomic sequencing level, and thus determine which other genes could alter its expression, possibly in a compensatory manner, in particular genes associated with efflux pumps or genes affecting bacterial fitness^54,55^. Likewise, the vulnerability of *MSMEG_0250* could be analyzed through a *knock-out* by homologous recombination, generating the most extreme scenario associated with a complete absence of expression.

In conclusion, silencing of the *MSMEG_0250* gene results in a significant 5-fold decrease in POA efflux velocity in M. smegmatis, so this gene could be involved in active POA transport.

## ACKNOWLEDGEMENTS

We thank Dr. Jeremy Rocks for his generous support in providing us with the technology to perform the CRISPRi technique.

